# Deterministic Formulas and Procedures for Stochastic Trait Introgression Prediction

**DOI:** 10.1101/2024.04.01.587554

**Authors:** Temitayo Ajayi, Jason LaCombe, Güven Ince, Trevor Yeats

**Affiliations:** Nature Source Improved Plants, 95 Brown St, Ithaca, 14850, New York, United States

**Keywords:** Trait introgression, stochastic modeling, mathematical analysis, optimization, genomics

## Abstract

**Key message:** We derive formulas for the background noise during trait introgression programs and use these formulas to quickly predict noise for up to five future generations without using simulation.

Trait introgression is a common method for introducing valuable traits into breeding populations and inbred cultivars. The process involves recurrent backcrossing of a donor individual (and its descendants) with a desirable, inbred line that lacks the aforementioned traits. The process typically concludes with a final generation of selfing in order to recover lines with the traits of interest fixed in the homozygous state. The particular breeding scheme is usually designed to maximize the genetic similarity of the converted lines to the recurrent parent while minimizing a breeders’ cost and time to recovering the near isogenic lines. Thus, key variables include the number of generations, number of crosses, and how to apply genotyping and selection during the process. In this paper, we derive analytical formulas that characterize the stochastic nature of residual donor geneome (i.e., “background noise”) during trait introgression. We use these formulas to predict the background noise in simulated trait introgression programs for five generations of progeny, as well as to construct a novel mathematical program to optimally allocate progeny to available parents. This provides a framework for the design of optimal breeding schemes for trait introgression involving one or more traits subject to the requirements of specific crops and breeding programs.

## 1 Introduction

The challenge of developing new material in response to increasing market pressure— need for improved yield for food and energy, resistance to disease and changing climate and microbial interactions—is ever present in plant breeding [1–3]. Typically, elite germplasm is enriched for favorable alleles for critical quantitative traits like yield but may lack one or more qualitative traits of interest that exist in non-elite germplasm. Trait introgression (TI) is an operation tool used to rapidly introduce desired traits into elite germplasm through backcrossing that results in one or more near isogenic lines (NILs) with high similarity to the elite (recurrent) parent while incorporating specific traits from a different donor line. This approach has been used to integrate increased performance for a variety of traits with different effects [4–7] that are conferred by native and/or transgenes [8]. Common examples include yield [4], disease resistance [9], drought tolerance [10], salinity tolerance [5], flood tolerance [11], herbicide tolerance [12], and maturity [13].

Many programs include TI as a core component of product/cultivar development strategy [14]. Material developed through breeding and pre-breeding efforts is inserted with the desired genes typically through multiple rounds of crossing. The introgression process is tracked with either phenotypes or molecular markers to ensure development of the desired end-product. Successful execution of the process involves the confirmation of the presence of the gene or phenotypic trait of interest within the developed material.

Common considerations for TI-execution include: the number of backcross and selfing generations, the use of phenotypic and/or molecular markers and the density of these markers, the target percentage of retained donor content (i.e., background noise), the total number of progeny to be produced in each generation, and the number of material advanced per generation. Challenges to these considerations include genotyping costs, greenhouse or field space limitations, time limitations, and seed-availability or productivity constraints inherent to the material.

Operations research (OR) has long been a source of solutions for optimal decision-making in the presence of various constraints. The increased availability of both molecular, phenotypic, and operational data within breeding programs, coupled with the continued development of computational power allows for improved analysis and understanding of the observed factors. Existing OR methodology for estimating or predicting the cost or benefit of TI strategies often involves the simulation and evaluation of mid- and end-point progeny (e.g., [15–19]), especially when some form of informed selection is involved in the proposed introgression scheme. Although simulation provides an effective approach, it can be computationally expensive for the sensitivity analysis required for the consideration of multiple strategies, especially when multiple TI projects are in execution or when evaluating the efficiency of new TI projects.

The proposed approach replaces simulation-based evaluation with deterministic calculations of the stochastic procedure using techniques from extreme value theory to model response to intermediate selection. We compare the approach with the results from simulation-based evaluation to demonstrate the speed and efficiency gains, as well as the additional tradeoff/sensitivity analyses and improved decision-making made possible by this approach.

There has been some related work in the cross-section between plant breeding and operations research. Studies in the marker-assisted selection space (e.g., [20–22]) provide context for where our work fits in the TI analysis literature. Most notably, [22] study both foreground and background selection in TI from a probabilistic- and simulation-based perspective. The authors derive the probability of successful offspring relative to the foreground, compute the necessary generation population sizes to produce successfully introgressed offspring, and also compute the error rates for the flanking markers’ representation of the quantitative trait locus (QTL). To build on the work of previous studies, we present our own novel approach in which we use a Poisson model to govern crossover events, derive specific formulas for the cases in which several transgenes are on the same chromosome, and consider the effects of selection pressure.

## 2 Methods

### 2.1 Preliminaries

We focus on successful trait introgression as measured by future progeny with: (1) all desired traits from the donor, and (2) a large percent of traits from the recurrent parent. The desired traits from the donor, along with nearby flanking markers, constitute the *foreground* of the genome, and the rest of the genome is the *background*.

For simplicity, we consider only diploids with a finite set of chromosome pairs *𝒞* (we refer to the chromosome pair as a “chromosome”). The recurrent parent and the donor individual are assumed to be homozygous. Thus, without loss of generality, we assume that there is a single recurrent parent that is homozygous for A alleles, and that there is a single donor individual that is homozygous for B alleles at any particular locus. We justify this generalization because if the recurrent parent and donor have the same genotype for some marker, that marker will remain the same throughout the breeding process. Thus, the set of markers in the genome, *ℳ*, consists only of markers in which the recurrent parent and donor differ (i.e., “free markers”). For each chromosome *c* ∈ 𝒞, let the marker set be given by 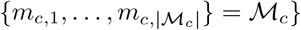, where the markers in the chromosome are indexed from “left to right,” i.e., from one end of the chromosome to the other.

The stochastic nature of the genetics depends on where the markers lie on the genome in relation to each other, including markers that will be limited to a fixed genotype. In the following, *m*_*j*_ indicates a free marker. We use *B*_*L*_(*m*_*j*_) or *B*_*L*_(*m*_*j*_, *m*_*k*_) to denote the right-most fixed marker that is to the left of *m*_*j*_ or to the left of both *m*_*j*_ and *m*_*k*_, respectively. We use *B*_*R*_(*m*_*j*_) or *B*_*R*_(*m*_*j*_, *m*_*k*_) to denote the left-most fixed marker that is to the right of *m*_*j*_ or to the right of both *m*_*j*_ and *m*_*k*_, respectively. When there is no such eligible fixed left marker, *B*_*L*_ is “defined” as a marker at -1000 cM, and similarly, when there is no eligible fixed right marker, *B*_*R*_ is “defined” as a marker at 1000 cM. In addition, we denote a fixed marker in between *m*_*j*_ and *m*_*k*_ by *B*_*I*_ (*m*_*j*_, *m*_*k*_), and we remark that there may be multiple such markers. Note that other markers may appear between these objects. We let *δ*(*m*_*j*_, *m*_*k*_) be the distance (in cM) between markers *m*_*j*_ and *m*_*k*_ on the same chromosome.

### 2.2 Foreground Analysis

The foreground of the genome is a specified area near the transgenes. We consider two types of foregrounds separately, each with its own application-driven use case.

The first type of foreground arises from when the TI gene is easily observable during phenotyping, or the TI gene can be directly genotyped with a genic molecular marker. For example, if there is a unique, dominant gene that confers resistance to an herbicide, then a simple test to determine resistance also reveals whether or not the individual has the relevant allele of the gene. Moreover, within the backcrossing portion of a trait introgression scheme concerning an herbicide resistance gene, it is immediately known—barring a rare mutation—that the individual is heterozygous at that locus. Thus, we do not need to track any nearby flanking markers, and the foreground is simply the marker of the transgene itself.

The second type of foreground comes from situations in which there is not an easy way to determine the genotype of the TI gene. For example, the TI gene’s approximate genetic position is known from association or QTL studies, but a genic marker is not available. In this case, the genotype of the TI gene must be inferred from genome-wide marker data. Thus, we use the nearby flanking markers as the foreground.

The accuracy of the flanking markers’ representation of the genotype at the TI gene depends on the distance from the TI gene to the associated flanking markers:

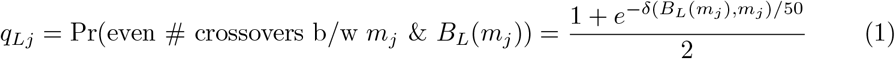

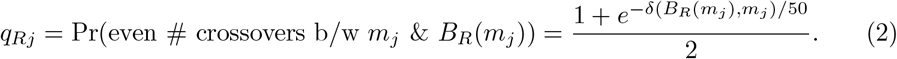

We can compute the conditional probability that the TI gene marker has the donor allele, given that the flanking markers have the donor allele

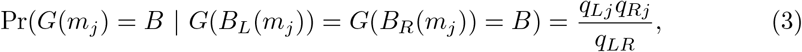

where *G*(*m*_*j*_) indicates the genotype of marker *m*_*j*_ and

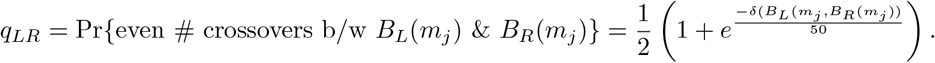

Note that *q*_*LR*_ = *q*_*Lj*_*q*_*Rj*_ + (1 *− q*_*Lj*_)(1 *− q*_*Rj*_).

Conditional probability is necessary because the offspring under consideration are only retained if the flanking markers have the donor allele.

From Equation 3, one can readily show with algebra

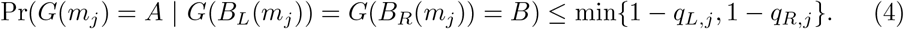

Thus, as long as one of the flanking markers is close to the position of the gene, the flanking markers provide a good indicator of the behavior of the gene, although a close flanking marker on both sides is multiplicatively better. Figure 1 shows that the error rate is generally low when both flanking markers are within 10 cM of the gene.

**Fig. 1:**
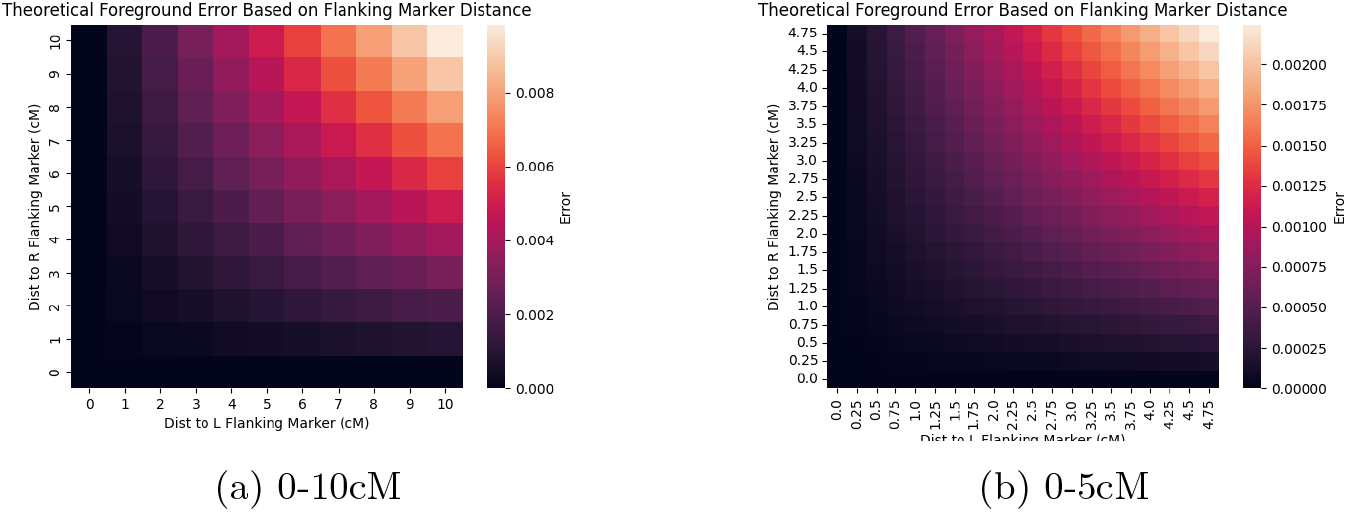
The probability of a progeny receiving no donor allele at the gene when both the immediate flanking markers receive donor alleles.

Whether the TI gene traits or alleles are easily observable, or the flanking markers have to be genotyped as proxies, within our context of trait introgression, one can only consider progeny that have all of the required TI genes (or proxies) for selection and advancement. For each chromosome *c*, let *𝒯*_1,*c*_ be the set of gene markers on chromosome *c* for traits that are phenotypically observable. Let *𝒯*_2,*c*_ be the set of flanking markers of genes on chromosome *c* for traits that are not phenotypically observable, where we assume that there is no overlap in flanking markers nor gene markers (i.e., every marker in *𝒯*_1,*c*_ ⋃ *𝒯*_2,*c*_ is unique). Finally, let 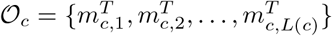 be the set of markers in *𝒯*_1,*c*_ ⋃ *𝒯*_2,*c*_ ordered according to their genetic position on chromosome *c*.

Suppose a progeny is produced from the cross of the recurrent parent and a donor that is heterozygous for all of the gene markers and their proxies, and is descended via backcrossing from the original donor individual. The probability that the progeny is also heterozygous for all of the gene markers and their proxies is

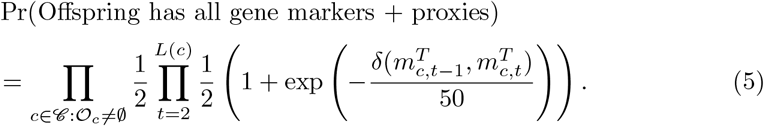

In general, the more genes involved, the lower the probability that a given progeny will possess all of the traits; hence, the choice of genes is a vital component of trait introgression. In the following, however, we assume that the genes have already been chosen with careful consideration.

### 2.3 Formulations for Stochasticity in Trait Introgression

In addition to using simulations to estimate the distribution of characteristics in future breeding generations, one can derive formulas towards the same aim. In this section we derive various formulas and methods to estimate the expectation and variance of the background noise of trait introgression progeny, based on the structure of the genetic map and the positions of the transgenes and the flanking markers. We verify these formulas and methods using simulation in Section 3.

There are three scenarios (eliminating symmetrically identical scenarios) to arrange fixed marker blocks and a single marker, which are illustrated in Figure 2. Also, there are scenarios in which a pair of markers can be oriented on a chromosome relative to blocks of fixed-allele markers, which are illustrated in Figure 3.

**Fig. 2:**
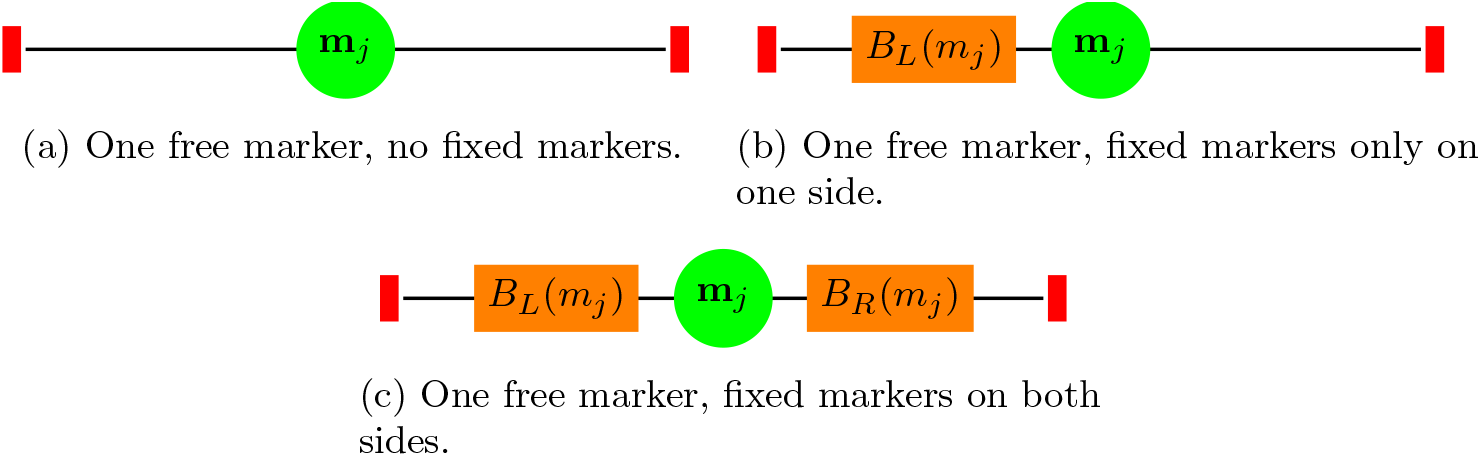
Single-marker scenarios

**Fig. 3:**
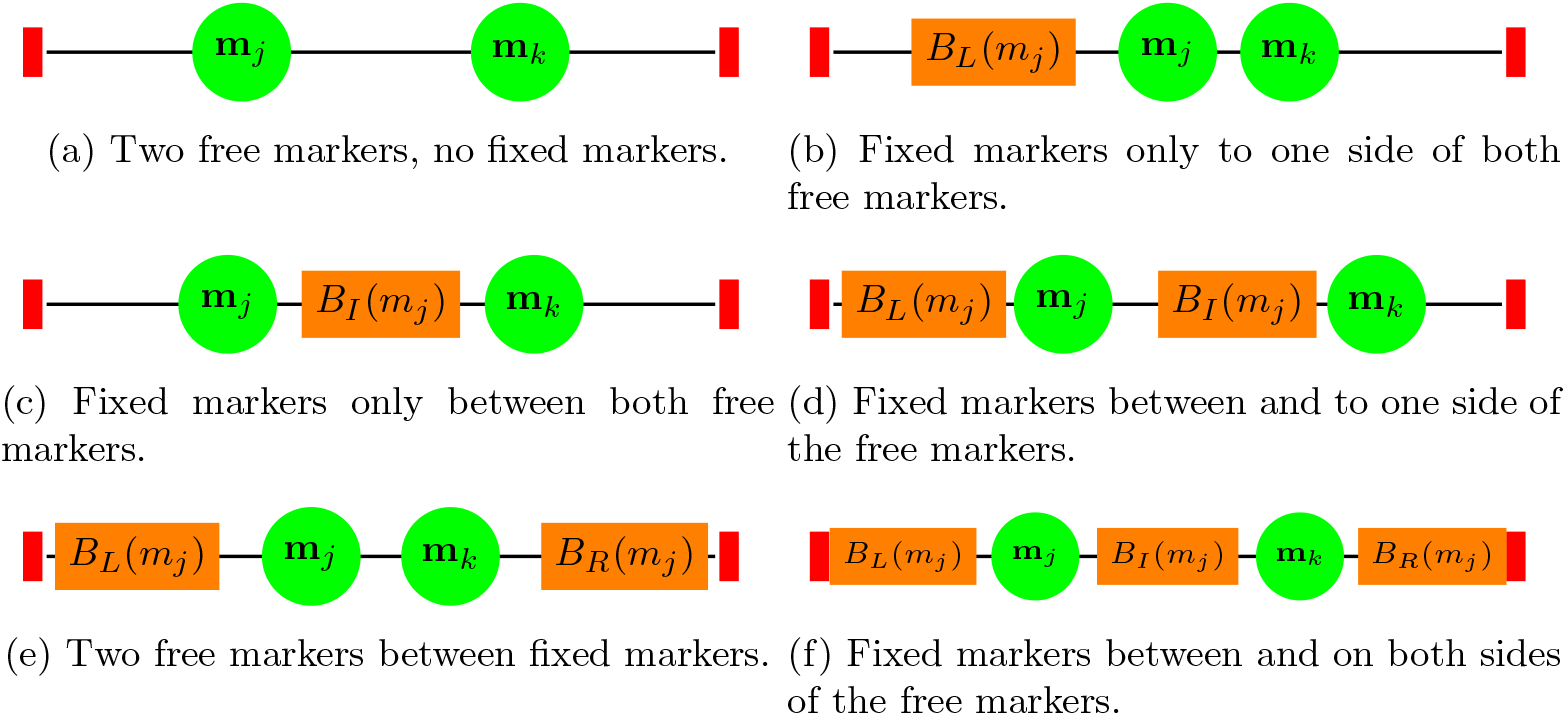
Two-marker scenarios

Although it is instructive to list all possible one- and two-marker scenarios, we can collapse these lists into a single scenario each. We use *X*_*j*_ to represent the Bernoulli random variable equal to 1 with the probability that the BC1 receives a B allele from the donor at marker *m*_*j*_. Also, the probability of an even number of crossovers between two free markers on the same chromosome *m*_*j*_ and *m*_*k*_ is *q*_*jk*_. Similarly, the probability of an even number of crossovers between a free marker *m*_*j*_ and a fixed marker *B*_*F*_ on the same chromosome is 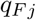; e.g., the probability of an even number of crossovers between *m*_*j*_ and *B*_*L*_(*m*_*j*_) is *q*_*Lj*_. The set of chromosome indices is *𝒞*. We use the expectation of *X*_*j*_ and the covariance between the random variables to derive the expectation and variance of the background noise.

To compute the expectation and variance of the background noise of progeny, it appears one must know, for each free marker, if there are fixed markers to the left (and if so, how close) and likewise for the right side. This allows one to choose the correct scenario (from Figures 2-3), which adjusts the probability of even crossovers. However, we can simplify this process by assuming that every marker has a fixed marker on either side of it, which reduces the number of scenarios to one. We place a “phantom” fixed marker at a large distance *D* to the left of the true left end of the chromosome, and we do the same with the right end of the chromosome. For example, a marker on a chromosome with no fixed markers would be bordered by a phantom fixed marker on either side, and *q*_*L,j*_ *≈ q*_*R,j*_ *≈* .5.

The same technique can be used for the covariance. Moreover, we can ignore any pairs of markers that are not immediately between the same two fixed markers— phantom fixed markers count as fixed markers in this context. This strategy eases the implementation of a formula-based approach to estimate the expectation and variance of the background noise.

#### 2.3.1 Single-Generation Formulas

Equation 6 is a formula to compute the expectation of *X*_*j*_, i.e., the probability that a given BC1 offspring has a B allele at marker *j*:

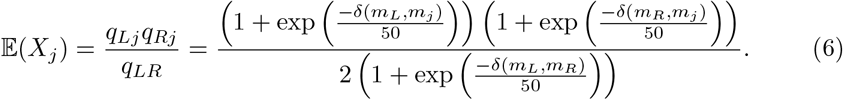

Thus, the formula for the expectation of the BC1 progeny background noise is

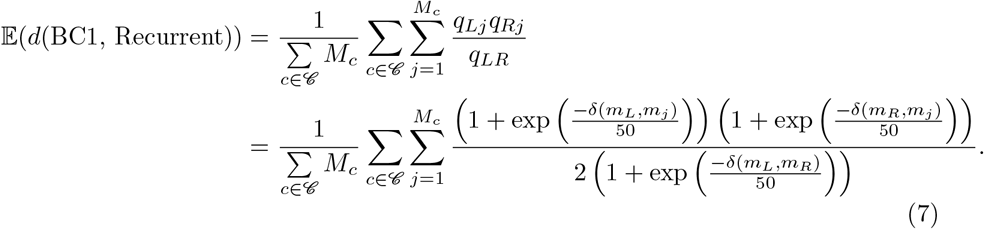

The formula for the covariance between marker *m*_*j*_ and marker *m*_*k*_ (assuming they are between the same two fixed markers, otherwise, it is 0) is given in Equation 8:

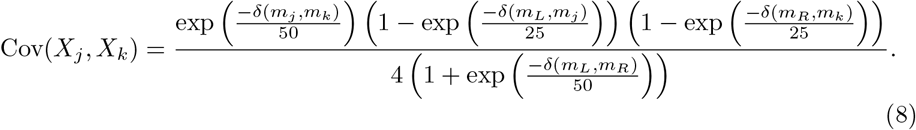

Hence, the formula for the variance of the background noise in the BC1 generation is

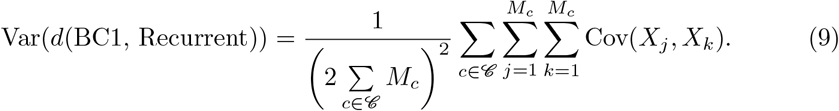

A derivation for (8) is available in C.1.

In addition to describing the distribution, from (6), one realizes that, given the positions of fixed markers, certain markers in the background are less likely to resolve and match the recurrent parent after meiosis. In fact, we show that the markers closest to the midpoint between the two immediately surrounding fixed markers are least likely to resolve.

##### Proposition 1.

*The probability that a marker, between two fixed markers, has a B genotype in a gamete, given that the progenitor is heterozygous at that marker, decreases as the cM distance to the midpoint between the fixed markers decreases*.

A proof of Proposition 1 is in the appendix.

### 2.4 Multi-Generation With Selection Formulas

Predicting background noise more than one generation into the future introduces one critical difficulty: the status of the markers in the non-elite parent are unknown. That is, we do not know with certainty if the parent is heterozygous or homozygous at marker *j*. In contrast, we know that the non-elite parent of the first generation (the F1) is heterozygous at marker *j*.

To account for the increased stochasticity, we generalize our single-generation prediction formulas. First, we observe that if we are crossing a generation *g* individual to make a generation *g* + 1 individual, the outcome at a marker *j* is a random variable 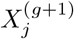 that is the product of two random variables, 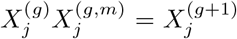. The first factor, 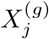 is the random variable that the generation *g* individual is heterozygous at marker *j*. The second factor, 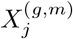, is the random variable that describes if the meiosis process for generation *g* leads to an even number of crossovers between *B*_*L*_(*m*_*j*_) and *m*_*j*_ and then also between *m*_*j*_ and *B*_*R*_(*m*_*j*_). Note that 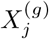 and 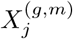 are independent of each other; the number of crossovers during meiosis is independent of the parent’s heterozygosity status. The meiosis random variable for marker *j* is binary where *h*_*j*_ is the probability of that that random variable equals 1:

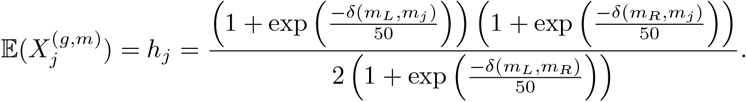

We also require the covariance of two meiosis random variables that are between the same two fixed markers (*η*_*j,k*_):

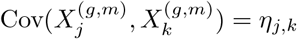

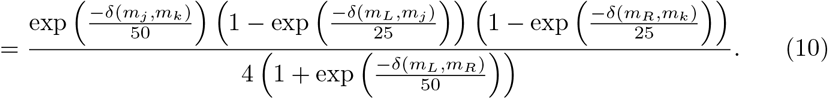

Note that *h*_*j*_ and *η*_*j,k*_ are the same in every generation.

Next, we consider the random variable that corresponds to the status of the markers of the parent, 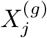. The mean and covariance of these variables, conditioned on the background noise, is given by the following quantities

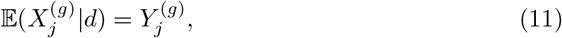

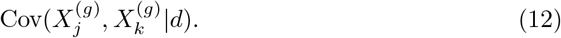

In the F1 generation (generation 0), the above quantities are known

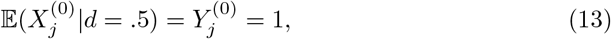

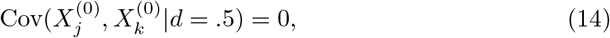

which makes it possible to compute these quantities for the next generation and estimate the effect of conditioning on background noise, and we will arrive at these steps shortly.

Given that 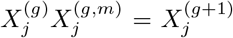 and that 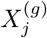 and 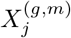 are independent, we can compute the unconditional expectation and covariances associated with 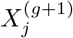.

#### Proposition 2.

*The expectation of* 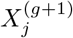 *is*

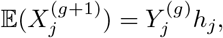

*and the covariance between* 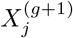 *and* 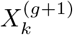, *when m*_*j*_ *and m*_*k*_ *are between the same consecutive fixed marker blocks, is*

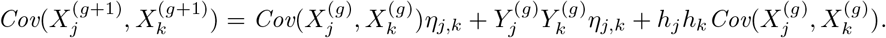

A proof of Proposition 2 is in the appendix. Again, when two markers are not between the same consecutive fixed markers, their covariance is 0.

#### Corollary 1.

*The mean and variance of the progeny background noise is given by*

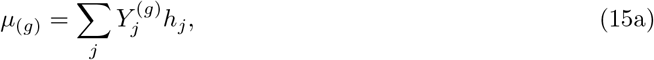

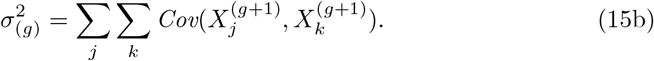

Given Corollary 1, we can analytically produce progeny for a given parent. Consider the *p*^*th*^-ranked parent of generation *g*; using Corollary 1, the mean and variance of its progenies’ background noise is *μ*_(*g,p*)_ and 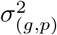, respectively, where *p* reflects the parent’s rank in generation *g*. Then, using a formula for normal distribution order statistics from [23], we can produce progeny with background noises 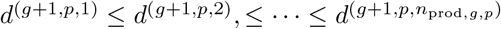:

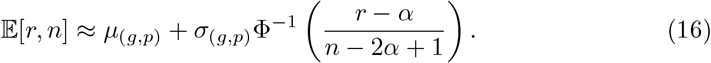

In (16), *r* is the rank of the progeny from a sample of *n* progeny, and from [23], we set the parameter *α* = *π/*8. We remark that we use the normal distribution even though there are some cases in which the normal distribution is not a good modeling choice (such as some extreme examples in which a chromosome only has heterozygous markers in a small, high density region).

The expectations and covariances in Proposition 2 enable us to produce the progeny in the next generation, but to estimate the marker statuses of the realized progeny, we need to account for the background noise as a condition. Conditioning the expectation and covariances based on some background noise *d* is more involved and would possibly slow down the prediction process. We use heuristics to estimate the effect of background noise conditioning.

For the expectation,

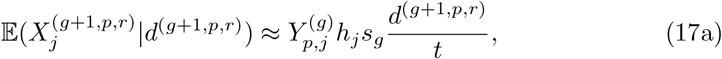

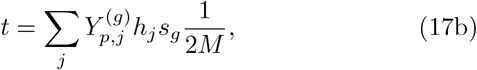

where *s*_*g*_ = 1 if generation *g* + 1 is formed from backcrossing and *s*_*g*_ = 2 otherwise and *p* is the index of the parent in generation *g*. We denote 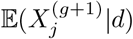 as 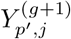, where *p′* is determined by that individual’s background noise ranking amongst all generation *g* + 1 individuals (from all parents). Note that (17) leads to 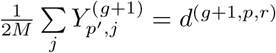.

For the covariance,

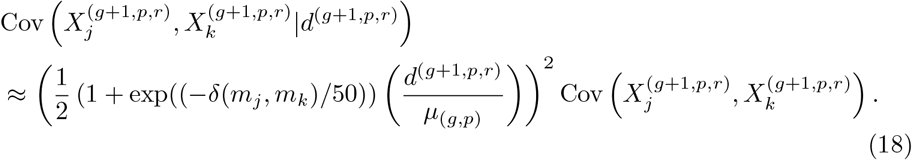

The factor with the squared ratio is a way to account for the mean of the random variable changing from *μ*_(*g,p*)_ to *d*^(*g*+1,*p,r*)^ after conditioning on the realized background noise. The factor with the exponential term makes the covariance decay faster for markers further apart.

We then select the *n*_*g*+1,*sel*_-best progeny to progress to the next generation as parents. These individuals are ordered by their background noise from [23]’s formula.

At this point, we have the expected marker statuses and marker status covariances for each parent in the *g* + 1 generation, and we can produce the *g* + 2 generation by restarting from the top of this section.

### 2.5 A Procedure for Fast, Analytical Estimation of Background Noise

In this section, we provide details on our implementation of a novel, efficient, simulation-free approach to estimating the background noise during a trait introgression breeding scheme over several generations. We use the formulas derived and presented in Sections 2.3-2.4 to produce estimates of important quantities at various stages in the process. In particular, the process we describe is for multiple-generation trait introgression with selection. We permit both backcrossing and selfing in the breeding scheme. The breeding scheme can be a single stage, or multiple generations with selection.

The inputs to the procedure are: a genetic map, the map position of each TI gene, or in lieu of the marker’s location, the location of the left and right flanking markers, the breeding scheme (how many progeny to produce and select in each generation, and what type of cross). All expectations and variances refer to the background noise.

1. ***F1 Creation*** During the start of trait introgression, the F1 individual is produced by crossing the recurrent parent (homozygous for allele A) with a donor line (homozygous allele B).
2. ***Generation 1 Population Expectation, Variance*** 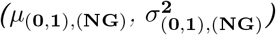 The F1 constitutes “Generation 0”. The progeny of this generation, which constitute Generation 1, has expectation (*μ*_(0,1),(*NG*)_) and variance 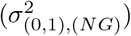 are estimated by 𝔼(*d*(BC1, Recurrent)) and Var(*d*(BC1, Recurrent)) from Equations 7 and 9, respectively, if the crossing type is backcross, or 2𝔼(*d*(BC1, Recurrent)) and 2Var(*d*(BC1, Recurrent)), if the crossing type is selfing.
3. ***Generation 1 Order Statistics (*d**^(**1**,**p**,**r**)^) The order statistics represent the ranking of background noise of the realized offspring in Generation 1, where *d*^(1,*p,r*)^ represents the *r*^*th*^-best (i.e., smallest) order statistic from the *p*^*th*^ family in the previous generation. In Generation 1, there is only one family (from the F1). The order statistic *d*^(1,1,*r*)^ is estimated from Equation 16 [23] using *μ*_(0,1),(*NG*)_ and 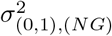, given the number of progeny produced per family in Generation 1 (*n*_0,prod_). Each selected progeny that progresses to be a parent has its marker statuses 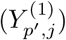 and covariances Cov 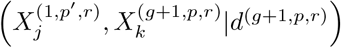 updated using Equations (17)-(18).
4. ***Generation 1 Selection (****𝒮*_**1**_) The *n*_1,sel_-best “progeny” from all available order statistics are selected as the parent set (*𝒮*_1_) for the next generation. In Generation 1, there is only one family; hence, *𝒮*_1_ is made up of the *n*_1,sel_-best offspring from the F1.
5. ***Next Generation Family Expectation, Variance*** 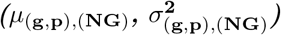 Given we are in generation *g*, selected individual *p* from *𝒮*_*g*_ will give rise to its family of progeny. We estimate *μ*_(*g,p*),(*NG*)_ and 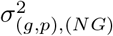 using Equation (15) if the crossing type is backcross or twice those values if the crossing type is selfing.
6. ***Generation* g** + **1 *Order Statistics*** The order statistics represent the ranking of the background noise realized offspring in Generation *g* + 1, where *d*^(*g*+1,*p,r*)^ represents the *r*^*th*^-best (i.e., smallest) order statistic from the *p*^*th*^ family in the previous generation (arising from the *p*^*th*^ parent in *𝒮*_*g*_). The order statistic *d*^(*g*+1,*p,r*)^ is estimated from Equation 16 [23] using *μ*_(*g,p*),(*NG*)_ and 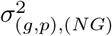, given the number of progeny produced per family (*n*_*g,prod*_) in Generation *g*.
7. ***Generation*** *g* + 1 ***Selection (****𝒮*_**g**_) The *n*_*g*+1,*sel*_-best “progeny” from all available order statistics are selected as the parents (*𝒮*_*g*+1_) for the next generation. Order the elements of 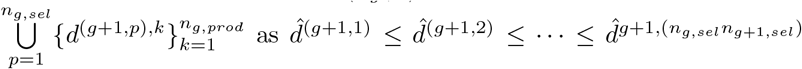. Then, *𝒮*_*g*+1_ consists of the parents with background noises 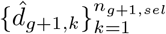.

Steps 4-7 are executed as many times as necessary to fulfill the breeding scheme. If one chooses, the parameters *n*_*g,prod*_ can be specified for different parent rankings (i.e., 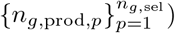, and we provide a method to do so online in Section 2.6.

### 2.6 Stochastic Optimization Model for Allocation

Suppose we have a set *ℐ* of eligible parents during the current generation *g* of the breeding program. As we have shown, based on the exact or predicted background noise, one can estimate the expectation, *μ*_*i*_, and variance, 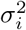 of the distribution of background noise in the progeny of parent *i*.. Thus, in the following, we use these estimates to construct an optimization model to best manage reproductive resources to decrease the background noise.

Let *𝒩* = {0, 1, …, *n*_max_} be the set of allowable progeny per eligible parent. Given *n* ∈ 𝒩, and *i* ∈ ℐ, the set of eligible parents, and *r* ∈ {1, …, *n*}, define the objective coefficient

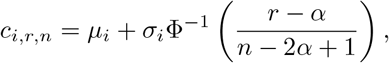

i.e., the expected *r*^*th*^ order statistic from parent *i* when the number of progeny is *n*. Also, let *S* be the number of progeny to select to proceed as eligible parents in the next generation. In the following model, the decision variable *x*_*i,r,n*_ indicates if the *r*^*th*^ progeny from the *i*^*th*^ parent (while producing *n* total progeny) is selected. The decision variable *y*_*i*_ represents the total number of progeny produced by parent *i*.

We now present the stochastic order statistic allocation model (SOSA):

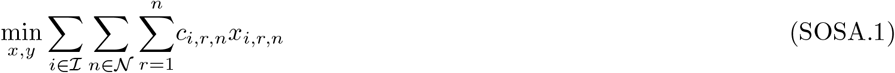

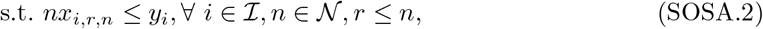

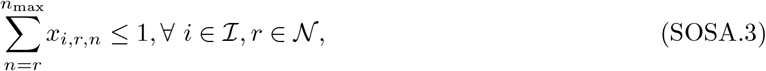

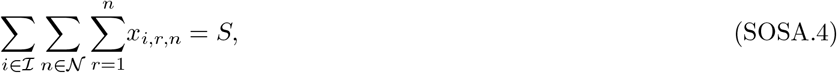

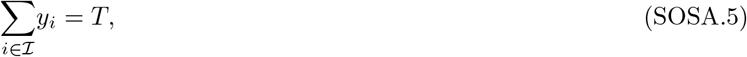

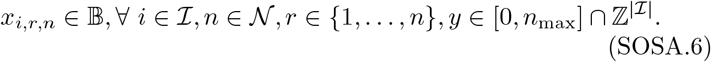

Here, *S* is the number of progeny to select and *T* is how many total progeny are allowed to be produced.

The objective function minimizes the sum of the expected order statistics of the selected progeny. Constraints (SOSA.2)-(SOSA.3), coupled with the objective, force the order statistics from a parent to come from a single sample size, dictated by *y*_*i*_. Constraint (SOSA.4) enforces that *S* total progeny are selected, and constraint (SOSA.5) ensures that *T* progeny are produced.

Given that the *x* decision variables are three-dimensional, one concern about SOSA may be the size of the model. However, the total number of binary variables is 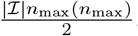. When | *ℐ*| = 20 and *n*_max_ = 20, the number of binary variables is 4200, which is typically a manageable number for a commercial solver, especially given that there are only 20 additional integer variables. In addition, there are 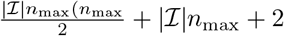 linear constraints.

One can include other constraints, such as a lower bound on the number of parents that contribute to the selected progeny, which may increase diversity, or parent-specific progeny capacities based off of seed projected seed availability.

## 3 Results

### 3.1 Single-Generation Backcrossing Predictions

We generated a homozygous for A allele lite individual, a homozygous B donor individual, and crossed these two individuals to produce an F1. Then, the F1 was backcrossed with the elite parent until 5000 BC1 offspring were produced with a B allele at each marker within the fixed marker blocks. This process was executed with the following parameters:

- *n*_Fixed_ ∈ {2, 4, 6, 8}, the number of transgenes that must be fixed in the backcrossed generations.
- *h* ∈ {.001, 2.5, 5, 10}, the half-length of the fixed marker block in cM. Each block emanates *h* cM to the left and right of the transgene at the center of it.

From the BC1 population, we computed the empirical expectation and variance of distance of BC1 individuals to the elite line. We then implemented the formulas in Section 2 to compute the variance of the number of B alleles in the backcrossed offspring, and converted that into the variance of the distance of the backcrossed progeny to the elite *among markers not in any fixed blocks*. This process was repeated for different randomly generated, ten-chromosome maps, quantities of selected transgenes, and different block lengths. Figure 4 shows the comparison of the variances computed empirically and by formula.

**Fig. 4:**
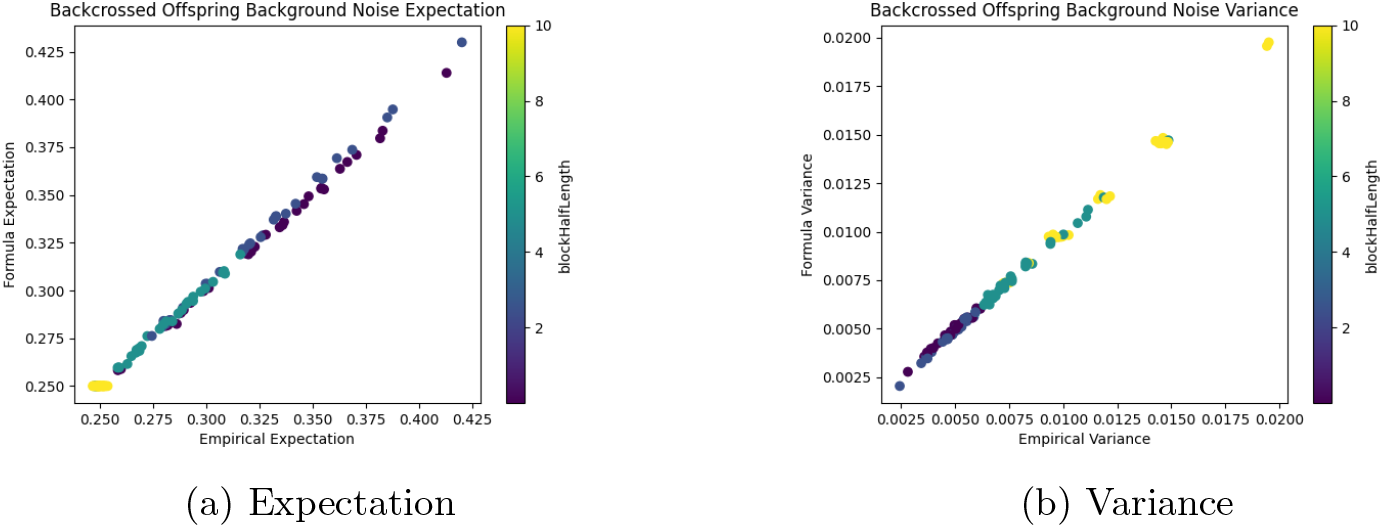
Comparison of empirical and formula-based expectation and variance of the distance between the BC1 generation and the elite line with color coding by block- HalfLength.

Overall, the formula matches the empirical result without any obvious signs of bias. From the color coding, we can see that as the length of the fixed marker block increases, there is less background noise but there is also more variance in the background noise.

### 3.2 Multi-Generation With Selection Predictions

In this section, we show the performance of our generalized, formulation-based procedure to predict the background noise during trait introgression. Various trait introgression settings were explored by using the following parameter values:

- *n*_Fixed_ ∈ {1, 2, 4, 6}, the number of transgenes that must be fixed in the backcrossed generations.
- *h* ∈ {.001, 5}, the half-length of the fixed marker block in cM. Each block emanates *h* cM to the left and right of the transgene at the center of it.
- Breeding Scheme (*n*_*sel*_, *n*_*prod*_, cross type) is one of the following:
  1. (10, 10, BC *×* BC *×* BC *×* BC *×* Self)
  2. (5, 10, BC *×* BC *×* BC *×* BC *×* Self)
  3. (5, 20, BC *×* BC *×* BC *×* BC *×* Self)
- The genome was generated randomly with one of the following three settings:
  1. 10 chromosomes of length 128 cM, with markers spaced by an average of 2.5 cM
  2. 10 chromosomes of length 168 cM, with markers spaced by an average of 5 cM
  3. 5 chromosomes of length 128, with markers spaced by an average of 1.25 cM.

For each set of unique parameter values, trait introgression was modeled for five independent trials with different randomly generated maps and random locations of the transgenes. To estimate the background noise at each generation empirically, we simulated the TI process 100 times for each trial.

Figure 5 shows the predictions for the expected background noise among non-fixed markers. Although the BC2-BC4 generations have a slight upward bias (see Figure 9 for the distribution of prediction error), the predictions are still close to the empirical values. The predictions for generation BC1 are the most accurate as they make use of the true expectation and variance values for a population formed by backcrossing an F1 and do not require approximations used in future generations.

**Fig. 5:**
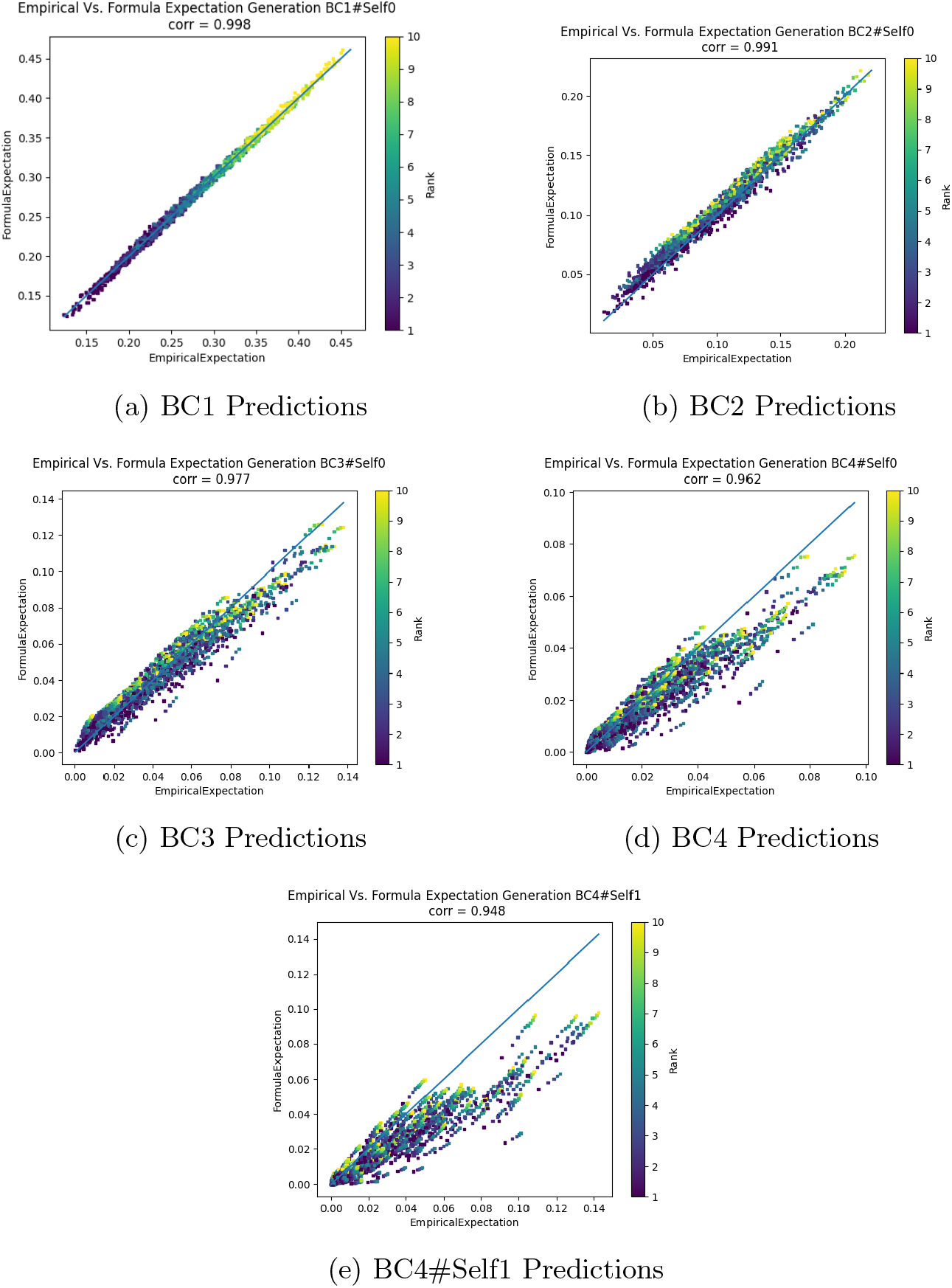
Predictions using formula-based procedure versus empirical estimates of the background noise for each generation during trait introgression. The color indicates the rank of the individual in the selection (e.g., the best BC2 has rank 1). Distributions of the error are available in the appendix.

Figure 6 depicts the predicted survivorship lines from progenitors of each generation. For instance, consider the breeding scheme which selects ten parents per generation, and each parent produces ten progeny (Figures 6a-6b). The blue line in Figure 6a indicates that in the first generation, the top four eligible parents nearly always have a progeny selected into the next generation, whereas those with a rank of 6 or greater rarely have offspring selected. The blue line in Figure 6b shows that a progeny from the top-ranked, first-generation parent exists in the final generation, whereas the fourth-ranked parent from the first generation, on average, only has a progeny in the second generation and rarely any descendants after that.

**Fig. 6:**
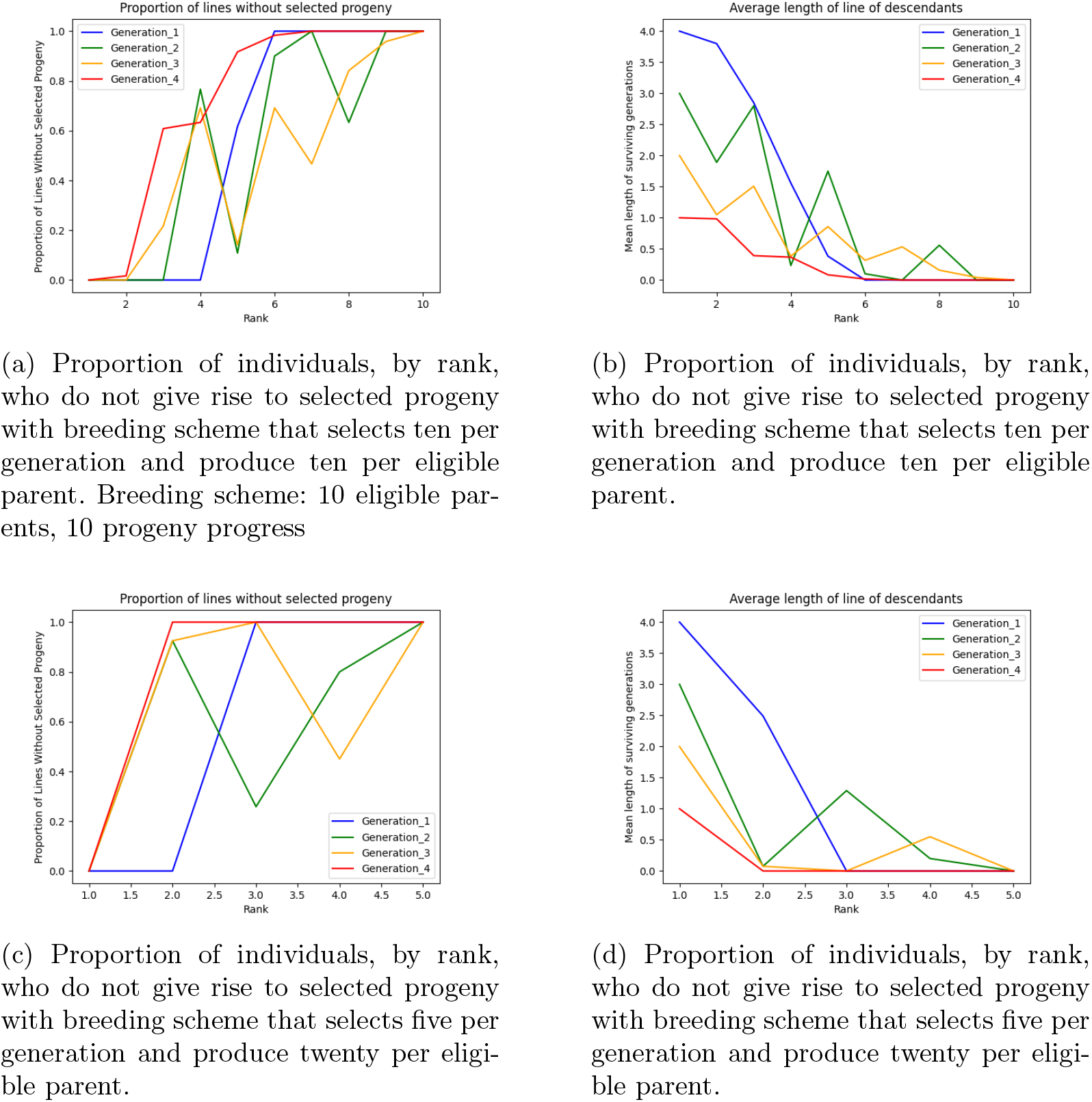
Lineage lengths through multiple breeding schemes.

Figure 7 shows the improvement in computational efficiency using our formula-based approach. From Figures 7a-7b, we observe that the main factor that increases computation time for the formula-based approach is the number of selected progeny, and the main factor that increases simulation time is the number of transgenes. The magnitude of relative improvement from simulation (on a natural logarithm scale) can be observed through Figure 7c. The simulation can take anywhere from 20 to a few thousand times longer than our formula-based approach.

**Fig. 7:**
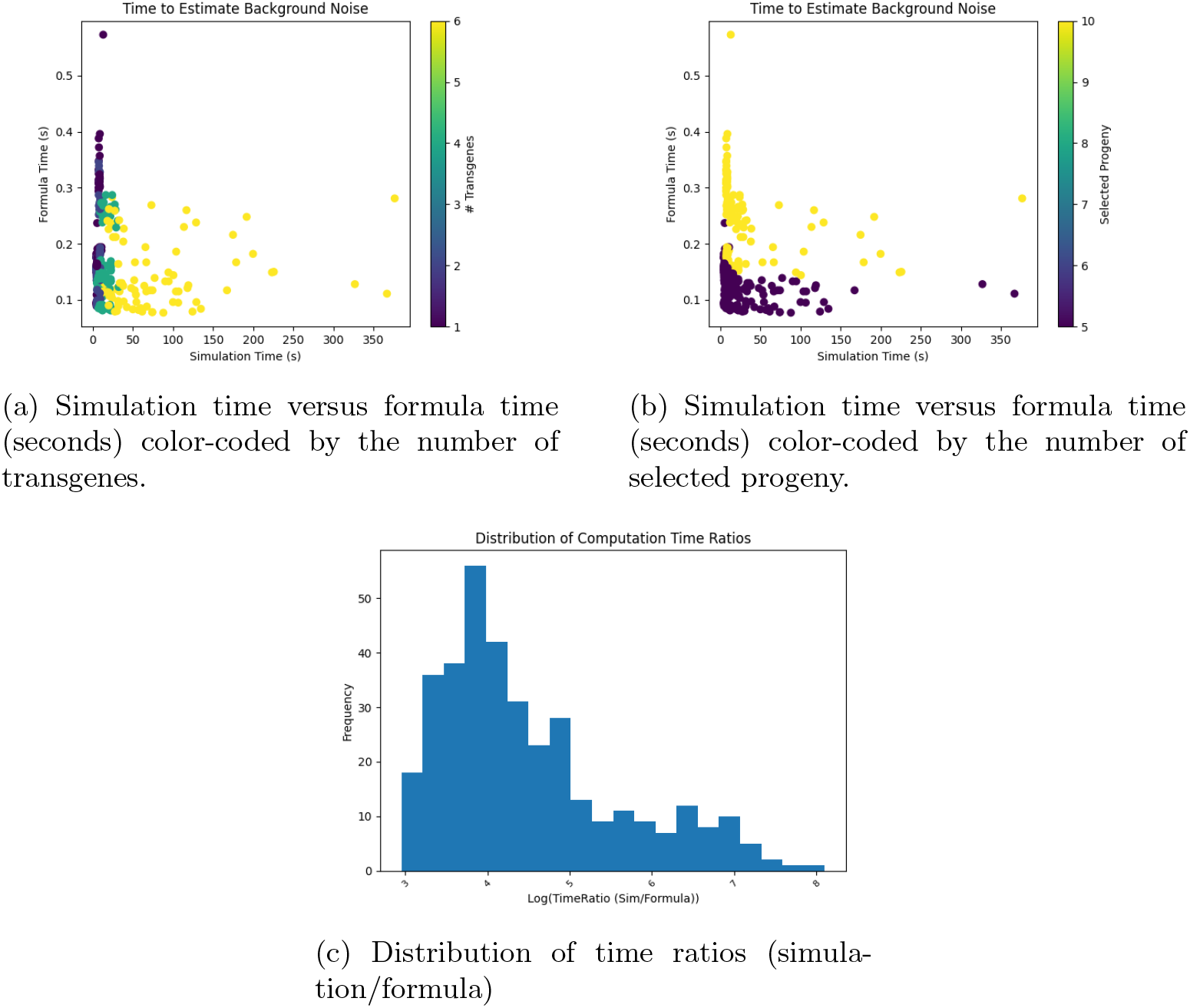
Comparison of computation time using simulation versus formula-based approach.

### 3.3 SOSA Results

To demonstrate the purpose of the SOSA model, we ran the optimization under various settings (see Table 1) using Gurobi 10.0.0 [24]. There are 84 unique parameter combinations, and each parameter combination was trialed with a unique, random genetic map 3 times. For each trial, we recorded the average expected background noise from the optimally selected progeny. Then, we repeated the optimization with a constraint that ensures at least six parents produce selected progeny, which may increase diversity at the cost of increased background noise.

**Table 1:**
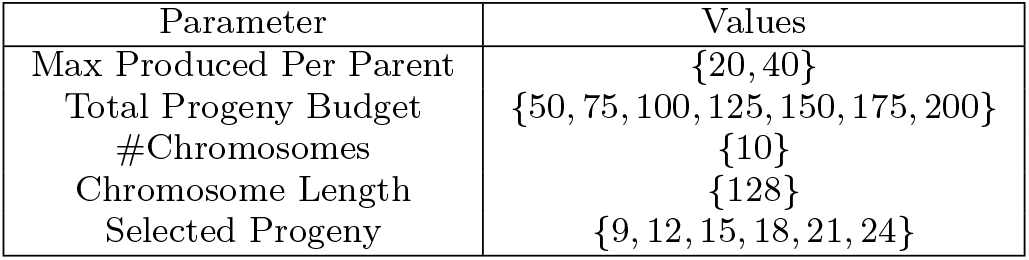
Parameters for SOSA Optimization trials.

Figure 8 shows the impact that both parent production capacity and as well as between the number of progeny to select have on the noise ratio (noise without additional constraint/noise with minimum parent constraint). As either parameter value decreases, the noise ratio decreases, which means the minimum parent constraint has a bigger effect.

**Fig. 8:**
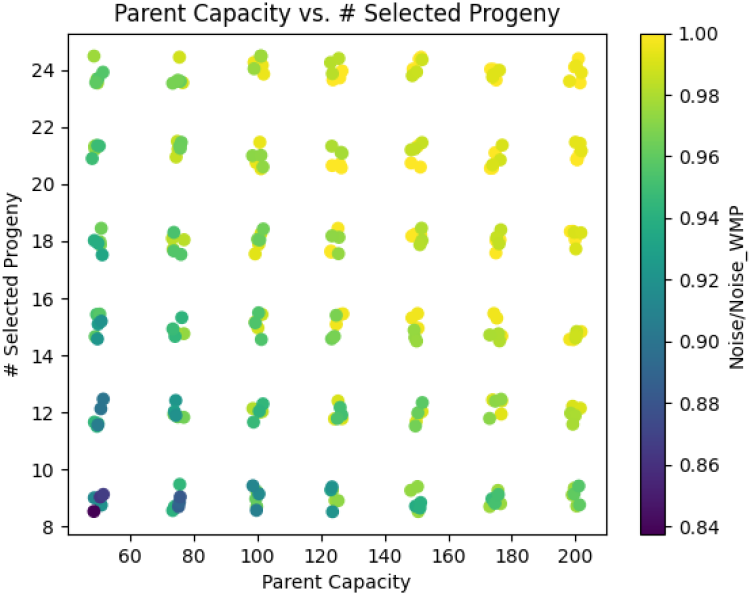
Production capacity per parent versus the total number of selected progeny, color-coded by the noise ratio (noise without minimum parent constraint divided by noise with minimum parent constraint). The parameter values were perturbed slightly in the plot to see more data points.

**Fig. 9:**
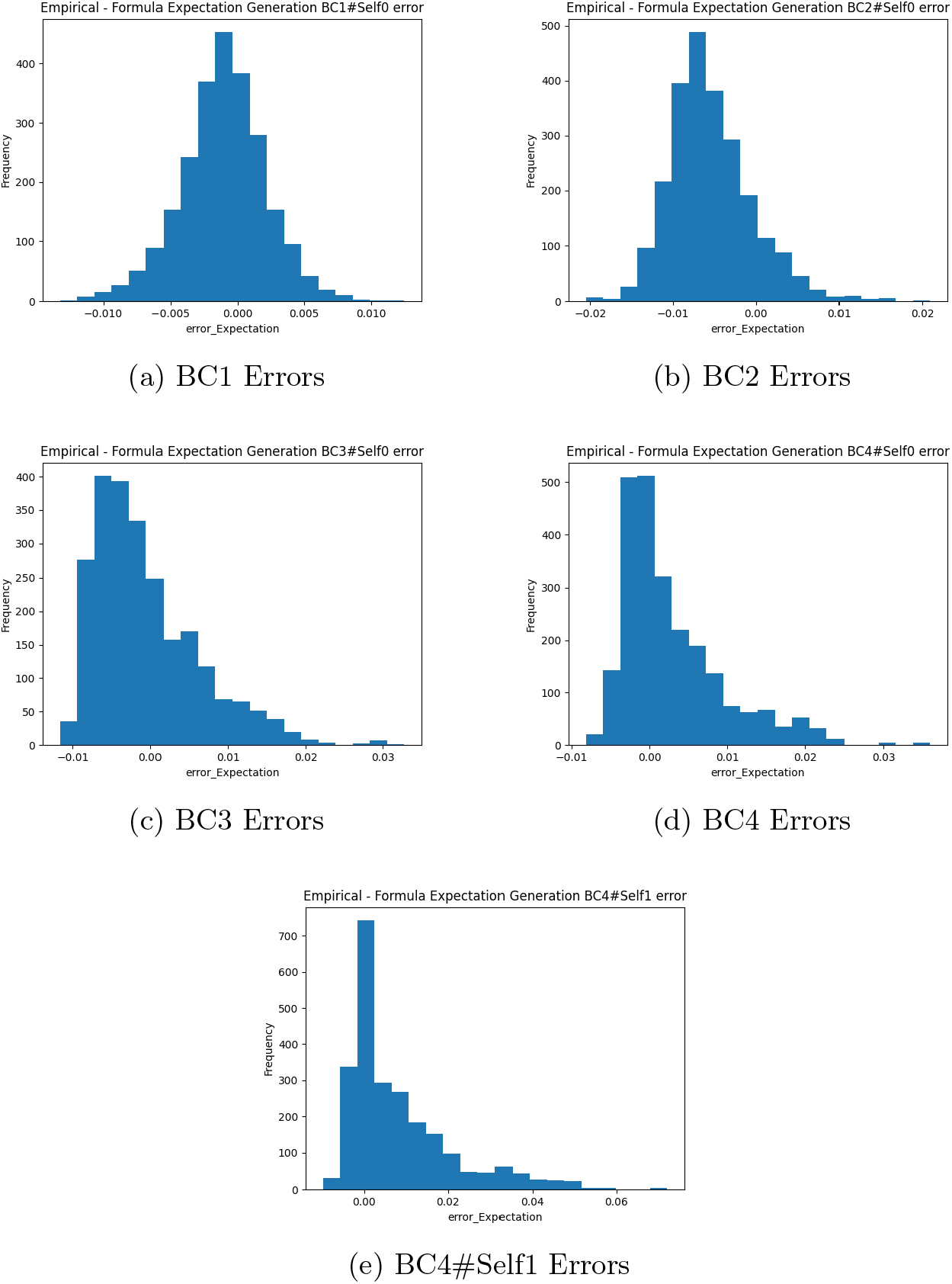
Errors for predictions using formula-based procedure versus empirical estimates of the background noise for each generation during trait introgression.

## 4 Discussion

The formula-based prediction scheme, while having limitations, is accurate at the relative level, albeit a bit less so at the absolute level. The predictions at the BC1 level are most accurate because these predictions are made with exact information; there are no heuristics to account for the inherent stochasticity of marker genotypes of future parents. This is not possible, to the best of our knowledge, when one wishes to predict background noise arbitrarily far into the future, so there is some decrease in accuracy in future generations. Notably, there appears to be a slight positive bias in the predictions of early generations. It should be noted that the “accuracy” is based on simulation results; nevertheless, there is a high correlation (greater than .94) after four backcrosses and selfing.

At the heart of our specific work in trait introgression prediction is the absence of simulation and the replacement with mathematical derivation and estimation. Simulation-based approaches (which we only use to validate our novel appraoch) can in various cases adequately aid prediction methods for complicated processes, such as trait introgression. However, the cost of simulation is the time needed to replicate the experiment or situation a sufficient number of times to achieve adequate convergence of the estimated quantities. In contrast, with our formula-based approach, computation time is minimal, and one does not need to repeat the computations to improve accuracy.

Further, this increased flexibility in our formula-based approach enables one to incorporate it into other complex decision-making problems. For example, one problem naturally linked to trait introgression is the allocation of progeny to eligible parents in the current generation (i.e., how many progeny should each parent introduce). In Section 2.6, we provide a Stochastic Order Statistic Allocation (SOSA) model that can be used to optimize progeny allocation, and its foundation is our formula-based prediction scheme. Although it is not included within this paper, due to brevity, the formulation-based predictions we provide can alternate steps with SOSA optimization solutions to predict progeny quality and also optimize progeny allocation within each generation.

In addition, although the accuracy of the predictions does decrease in future generations, the accuracy the predictions provide can help the design of a TI program. In a realistic scenario, genotyping may also occur during the intermediate stages (e.g., after a backcross). In this case, predictions can be updated based on the most recent genotype data available, which would likely result in even better accuracy.

There are several limitations of this study. First, part of our ability to use formula-based prediction methods relies on estimations of order statistics from a normal distribution. Although it appears to work well in the context of this study, we have not mathematically proven a justification for the use of the normal distribution. Second, our analysis of background noise is based on the fact that progeny are screened out immediately if they do not have the appropriate genotypes in the fixed regions. We note that this is the case in some trait introgression projects, including some in the industrial sphere, but not all programs fit this requirement.

One area for future research is coupling an optimization model, such as SOSA, with our formulation-based predictions. Although our predictions already include selection pressure, incoporating SOSA would help ease the burden of optimally designing a full breeding scheme. Another area of future research is applying our methodology to a real trait introgression program. On the theoretical side, two interesting questions include a mathematical justification for using the normal distribution for order statistic estimation and the source of the bias in predictions in later generations.

## Supplementary Information

This article has an accompanying appendix.

## Authors’ Contributions

All authors contributed to the study conception and design. Simulations and experiments were performed by TA and JL. Theoretical analysis and mathematical modeling was conducted by TA, JL, and GI. All authors contributed to the preparation and review of the drafts and final manuscript.

## Data and Code Availability

Currently, the code used to generate the data and execute the analysis consists of proprietary material of Nature Source Improved Plants. If necessary, upon request, the authors may reconstruct the code without the proprietary technology.

## Declarations

### Ethics approval

This is a mathematics- and simulation-based study. No ethics approval is relevant to this study.

### Consent to participate

There were no participants in this study.

### Consent for publication

There were no participants in this study.

## Appendix A Additional Figures

## Appendix B Omitted Proofs

### Proposition 1.

*Proof*. We derive the probability that a marker is *B* based on its position in between the two fixed markers, then take the derivative of that function twice to prove the claim.

Without loss of generality, let the left fixed marker be given position 0 and the right fixed marker *P*. Define the function *F* : [0, *P*] *→* [0, 1] as the probability that a marker at position *t* ∈ [0, *P*] has the genotype *B* in a gamete whose progenitor is heterozygous at that marker and the fixed markers, conditioned on the gamete receiving a *B* genotype for the fixed markers. Note that if a marker is at position *t*, then it is distance *t* to the left fixed marker and distance *P − t* to the right fixed marker. Thus,

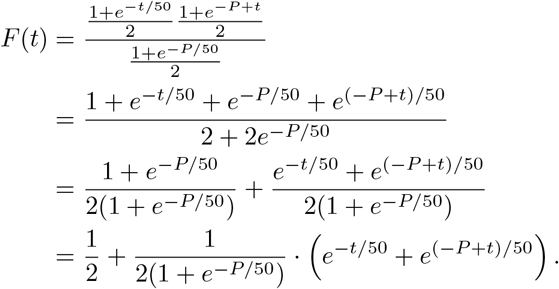

Taking the derivative with respect to *t*, we have

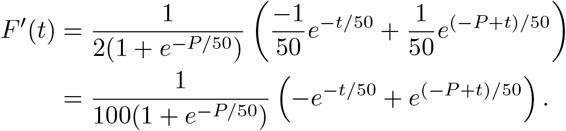

Solving for *F′*(*t*) = 0 gives *t*^*\**^ = *P/*2. Moreover, taking the second derivative, we have

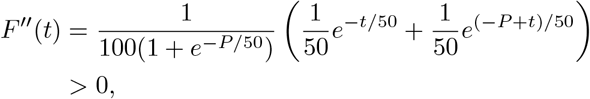

which implies that the unique value for which *F′* (*t*) = 0 is *t*^*\**^ = *P/*2, *F* is strictly convex, and *P/*2 is the unique minimum of *F*.

Therefore, the probability that a gamete receives a B genotype at a marker at position *t* is lowest at the midpoint, *P/*2 and the probability increases as one moves towards either fixed marker away from the midpoint.

### Proposition 2.

*The expectation of* 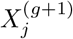 *is*

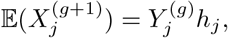

*and the covariance between* 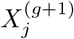 *and* 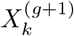, *when m*_*j*_ *and m*_*k*_ *are between the same consecutive fixed marker blocks, is*

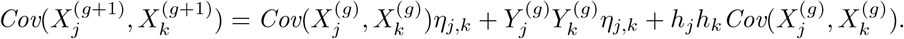

*Proof*. First, consider the expectation:

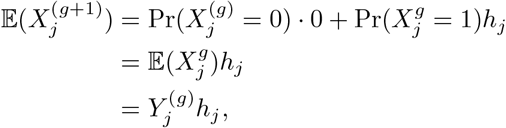

as claimed.

Next, consider the covariance. Observe that

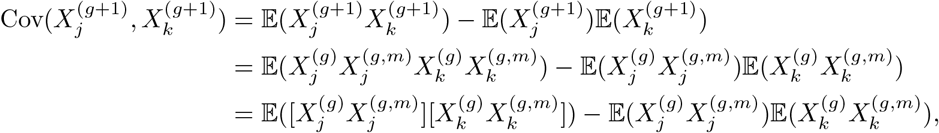

where we have so far, just used the definition of covariance along with 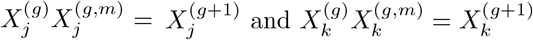.

Using the independence between 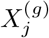 and 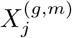 (and similarly, between 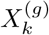 and 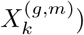, we have

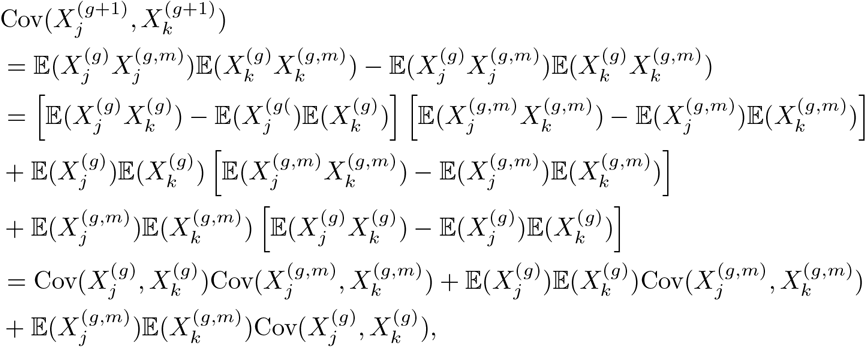

which proves the claim.

## Appendix C Omitted Derivations

### C.1 Single-Generation Covariance

In this section, we derive the equation

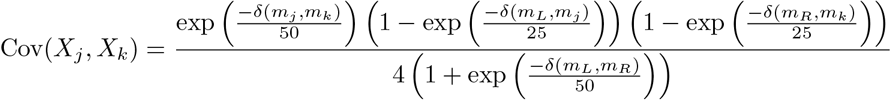

from (8). Observe, from the definition of covariance and using the even crossover probabilities (e.g., *q*_*Lj*_), that

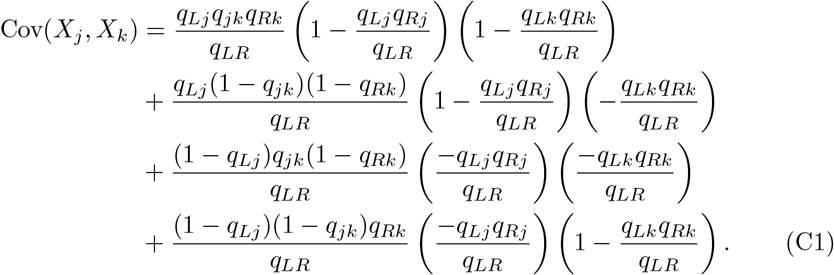

We then factor out 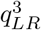 to obtain

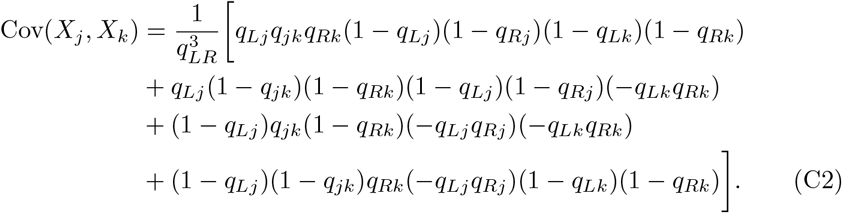

We subsequently factor out *q*_*Lj*_(1 *− q*_*Lj*_)*q*_*Rk*_(1 *− q*_*Rk*_) and expand terms to obtain

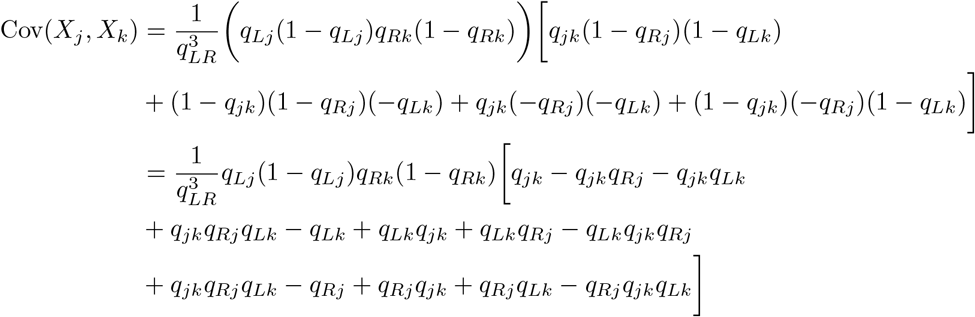

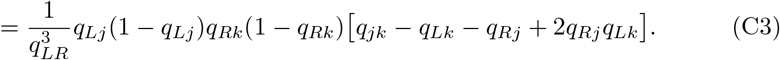

Observe that, by substituting for even crossover probabilities, we have

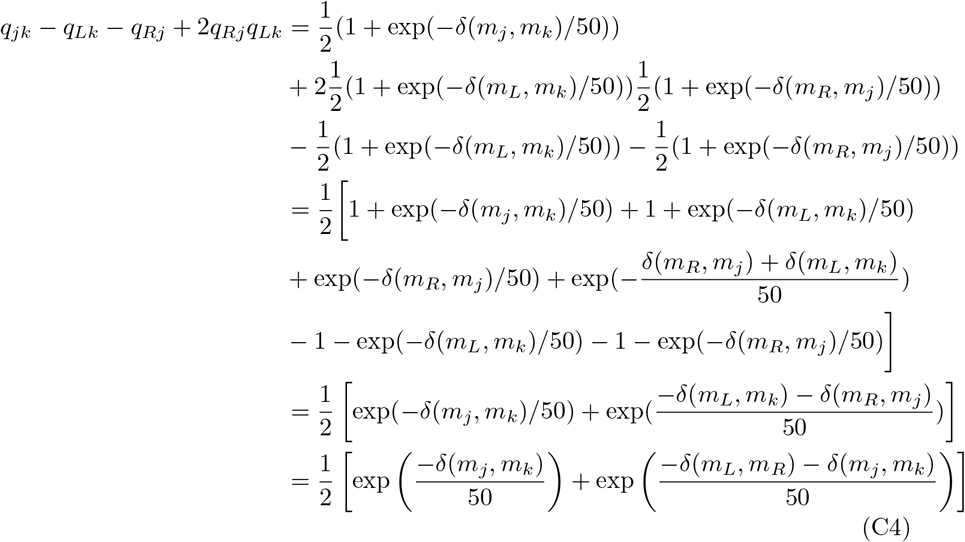

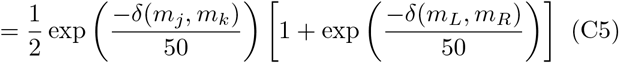

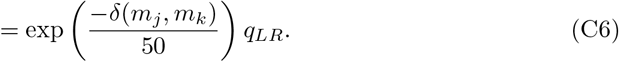

Hence, from substituting (C6) into (C3) and then substituting for even crossover possibilities, we have

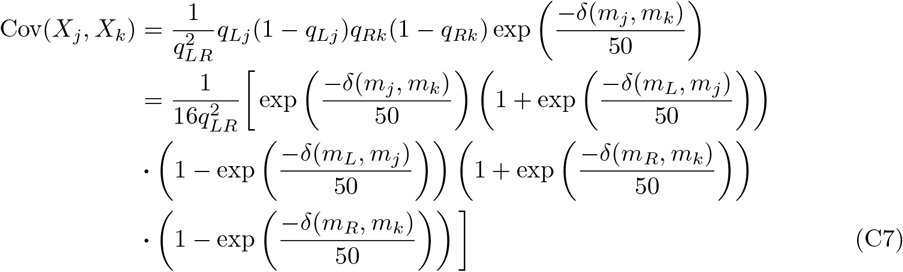

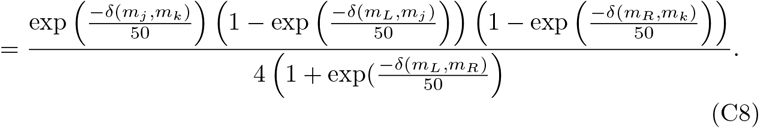

f

## References

[1] Araújo, S.S., Beebe, S., Crespi, M., Delbreil, B., González, E.M., Gruber, V., Lejeune-Henaut, I., Link, W., Monteros, M.J., Prats, E., et al.: Abiotic stress responses in legumes: strategies used to cope with environmental challenges. Critical Reviews in Plant Sciences 34(1-3), 237–280 (2015)

[2] Cheng, Y.T., Zhang, L., He, S.Y.: Plant-microbe interactions facing environmental challenge. Cell Host & Microbe 26(2), 183–192 (2019)

[3] Langridge, P., Braun, H., Hulke, B., Ober, E., Prasanna, B.: Breeding crops for climate resilience. Theoretical and Applied Genetics 134(6), 1607–1611 (2021)

[4] Concibido, V., La Vallee, B., Mclaird, P., Pineda, N., Meyer, J., Hummel, L., Yang, J., Wu, K., Delannay, X.: Introgression of a quantitative trait locus for yield from glycine soja into commercial soybean cultivars. Theoretical and Applied Genetics 106, 575–582 (2003)

[5] Qiu, X., Yuan, Z., Liu, H., Xiang, X., Yang, L., He, W., Du, B., Ye, G., Xu, J., Xing, D.: Identification of salt tolerance-improving quantitative trait loci alleles from a salt-susceptible rice breeding line by introgression breeding. Plant Breeding 134(6), 653–660 (2015)

[6] Sreeman, S.M., Vijayaraghavareddy, P., Sreevathsa, R., Rajendrareddy, S., Arakesh, S., Bharti, P., Dharmappa, P., Soolanayakanahally, R.: Introgression of physiological traits for a comprehensive improvement of drought adaptation in crop plants. Frontiers in Chemistry 6, 92 (2018)

[7] Antoine, A., Moreau, L., Charcosset, A., Teyssèdre, S., Lehermeier, C.: Usefulness criterion and post-selection parental contributions in multi-parental crosses: Application to polygenic trait introgression. G3-Genes Genomes Genetics 9, 3–4001292019 (2019)

[8] Gao, L., et al.: Do transgenesis and marker-assisted backcross breeding produce substantially equivalent plants? a comparative study of transgenic and backcross rice carrying bacterial blight resistant gene xa21. BMC Genomics 14 (2013)

[9] Hickey, L.T., Germán, S.E., Pereyra, S.A., Diaz, J.E., Ziems, L.A., Fowler, R.A., Platz, G.J., Franckowiak, J.D., Dieters, M.J.: Speed breeding for multiple disease resistance in barley. Euphytica 213, 1–14 (2017)

[10] Varshney, R.K., Gaur, P.M., Chamarthi, S.K., Krishnamurthy, L., Tripathi, S., Kashiwagi, J., Samineni, S., Singh, V.K., Thudi, M., Jaganathan, D.: Fast-track introgression of “qtl-hotspot” for root traits and other drought tolerance traits in jg 11, an elite and leading variety of chickpea. The Plant Genome 6(3), 2013–07 (2013)

[11] Breseghello, F., Coelho, A.S.G.: Traditional and modern plant breeding methods with examples in rice (oryza sativa l.). Journal of Agricultural and Food Chemistry 61(35), 8277–8286 (2013)

[12] Croser, J., Mao, D., Dron, N., Michelmore, S., McMurray, L., Preston, C., Bruce, D., Ogbonnaya, F.C., Ribalta, F.M., Hayes, J., Lichtenzveig, J., Erskine, W., Cullis, B., Sutton, T., Hobson, K.: Evidence for the application of emerging technologies to accelerate crop improvement – a collaborative pipeline to introgress herbicide tolerance into chickpea. Frontiers in Plant Science 12 (2021)

[13] Chandnani, R., Kim, C., Patel, H. JD amd Guo, Shehzad, T., Wallace, J., He, D., Zhang, Z., Adhikari, J., Khanal, S., Chee, P., Paterson, A.: Identification of small effect quantitative trait loci of plant architectural, flowering, and early maturity traits in reciprocal interspecific introgression population in cotton. Frontiers in Plant Science 13 (2022)

[14] Hernandez, J., Meints, B., Hayes, P.: Introgression breeding in barley: Perspectives and case studies. Frontiers in Plant Science 11, 761 (2020)

[15] Peng, T., Sun, X., Mumm, R.H.: Optimized breeding strategies for multiple trait integration: I. Minimizing linkage drag in single event introgression. Molecular Breeding 33(1), 89–104 (2014)

[16] Peng, T., Sun, X., Mumm, R.H.: Optimized breeding strategies for multiple trait integration: II. Process efficiency in event pyramiding and trait fixation. Molecular Breeding 33(1), 105–115 (2014)

[17] Han, Y., Cameron, J.N., Wang, L., Beavis, W.D.: The predicted cross value for genetic introgression of multiple alleles 205(4), 1409–1423 (2017)

[18] Han, Y., Cameron, J.N., Wang, L., Pham, H., Beavis, W.D.: Dynamic programming for resource allocation in multi-allelic trait introgression. Frontiers in Plant Science 12 (2021)

[19] Moeinizade, S., Han, Y., Pham, H., Hu, G., Wang, L.: A look-ahead monte carlo simulation method for improving parental selection in trait introgression. Scientific Reports 11(3918) (2021)

[20] Visscher, P.M., Haley, C.S., Thompson, R.: Marker-assisted introgression in backcross breeding programs 144, 1923–1932 (1996)

[21] Hospital, F., Moreau, L., Lacoudre, F., Charcosset, A., Gallais, A.: More on the efficiency of marker-assisted selection. Theoretical and Applied Genetics 95, 1181–1189 (1997)

[22] Hospital, F., Charcosset, A.: Marker-assisted introgression of quantitative trait loci. Genetics 147, 1469–1485 (1997)

[23] Blom, G.: Statistical estimates and transformed beta-variables. Phd thesis, Stockholm University, Stockholm, Sweden (1958)

[24] Gurobi Optimization, LLC: Gurobi Optimizer Reference Manual (2023). https://www.gurobi.com

